# Air-Liquid-Interface Reorganizes Membrane Lipid and Enhance Recruitment of Slc26a3 to Lipid-Rich Domains in Human Colon

**DOI:** 10.1101/2022.12.22.521645

**Authors:** Ming Tse, Yan Rong, Zixin Zhang, Ruxian Lin, Rafiq Sarker, Mark Donowitz, Varsha Singh

## Abstract

**Background and Aims:** Cholesterol-rich membrane domains, also called lipid rafts (LR), are specialized membrane domains that provide a platform for intracellular signal transduction. Membrane proteins often cluster in LR that further aggregate into larger platform-like structures that are enriched in ceramide and are called ceramide-rich platforms (CRPs). The role of CRPs in the regulation of intestinal epithelial functions remains unknown. Down Regulated in Adenoma (DRA) is an intestinal Cl^-^/HCO_3_^-^ antiporter which is enriched in LR. However, little is known regarding the mechanisms involved in the regulation of DRA activity.

**Methods:** Air liquid interface (ALI) was created by removing apical media for a specified number of days from 12-14 days post confluency Caco-2/BBe cells or confluent colonoid monolayer grown as submerged cultures. Confocal imaging was used to examine the dimensions of membrane microdomains that contain DRA.

**Results:** DRA expression and activity were enhanced by culturing Caco-2/BBe cells and human colonoids using an ALI culture method. ALI causes an increase in acid sphingomyelinase (ASMase) activity, an enzyme responsible for enhancing ceramide content in the plasma membrane. ALI cultures expressed a larger number of DRA-containing platforms with dimensions >2 μm compared to cells grown as submerged cultures. ASMase inhibitor, desipramine disrupted CRPs and reduced the ALI-induced increase in DRA expression in the apical membrane. Exposing normal human colonoid monolayers to ALI increased the ASMase activity and enhanced differentiation of colonoids along with enhancing basal and forskolin-stimulated DRA activity.

**Conclusions:** ALI increases DRA activity and expression by increasing ASMase activity and platform formation in Caco-2/BBe cells and by enhancing the differentiation of normal human colonoids.

**Synopsis:** Air-liquid interface (ALI) enhances total and brush border DRA expression and activity in Caco-2/BBe cells and human colonoids by causing differentiation of enterocytes and acid sphingomyelinase-dependent enhanced retention of DRA in ceramide-rich platform-like structures at the plasma membrane.

## Introduction

The intestinal brush border Cl^-^/HCO_3_^-^ exchanger DRA (SLC26A3) plays a major role in apical Cl^-^/HCO_3_^-^exchange in the human duodenum, ileum, and colon^1–4^. DRA takes part in both intestinal Cl^-^ absorption in which it is linked to the Na^+^/H^+^ exchanger 3 (NHE3), and HCO_3_^-^secretion where it is linked with CFTR and functions to reabsorb some of the Cl secreted by CFTR. Plasma membrane trafficking, lipid raft distribution, and protein-protein interactions are the currently identified pathways involved in the acute regulation of DRA activity in intestinal enterocytes. Reported studies have focused on the mechanisms involved in DRA inhibition in inflammatory diarrheal diseases, including IBDs, and that caused by Enteropathogenic *Escherichia coli*^5, 6^. However, very little is known about the factors involved in the activation of DRA activity and the regulation of its secretory function. In polarized enterocytes, DRA is expressed in the apical membrane (but also associated with the tight junctions) and in endosomal pools^7–9^. In the enterocyte brush border (BB), DRA resides as two distinct populations; one is diffusely distributed and the other is in lipid rafts (LR)^7^. Other intestinal BB ion transport proteins, including NHE3 and the chloride channel CFTR, are also in part distributed in BB LR^10–12^. The factors determining the distribution of proteins in the LR of well-differentiated intestinal epithelial cells are not well understood. Extensive studies recently have focused on the changes in CFTR activity that are associated with clustering in LR in pulmonary epithelial cells^13, 14^. While the activity of DRA is known to increase with its presence in LR^7^, little is known about the mechanisms regulating DRA localization in LR as well as the role of LR in the basal or regulated activity of DRA.

Lipid rafts are dynamic structures that are assembled from clustered lipids, principally, cholesterol and sphingomyelin (SM), that form a spatially and biochemically distinct domain that segregates specific membrane proteins within the plasma membrane and intracellular membranes. Plasma-membrane ceramides are often generated in lipid rafts upon the activation of the enzyme acid sphingomyelinase (ASMase), which cleaves SM into ceramide and a phosphorylcholine (PC)^15, 16^. Ceramide converted from SM in lipid rafts exerts its biological activity by altering the structure of lipid rafts^15^. Evidence from studies in airway epithelial cells has suggested ALI as a way to activate ASMase^1, 2^. In these studies, epithelial cells were typically seeded on the upper surface of semi-permeable membrane inserts, and cells were fed basolaterally with a culture medium. Contact with air on one side and nutrients on the other imposes polarity and partially mimics *in vivo* conditions of mature adult airways.

In the current study, we demonstrated the presence of raft-like clusters of DRA in the apical domain of Caco-2/BBE cells that were closely associated with acid sphingomyelinase (ASMase). In basal conditions, ASMase was bound to the extracellular surface of the BB of Caco-2/BBe cells. In the ALI cultures, the ASMase was activated and was present in aggregates along with aggregated DRA on the plasma membrane, including in detergent-insoluble fractions.

These were associated with enhanced DRA expression and increased basal and forskolin-stimulated DRA activity. Enhanced formation of platform-like structures is a well-established signaling mechanism under a wide range of pathological conditions^4, 14^. The present results indicate that these platform-like structures also contribute to the activation of DRA in intestinal epithelial cells. The acute increase in DRA activity due to the formation of BB ASMase-dependent platform-like structures is a newly defined mechanism of DRA regulation.

## Results

### ALI culture increases DRA expression at the apical membrane in Caco-2/BBe cells

To study the effect of ALI exposure, the apical media were removed from 12-14 days post confluency Caco-2/BBe cells were grown as submerged cultures, and the cells were studied 2 days later (**Fig 1A**). The 2-day ALI exposed cells were significantly taller compared to the cells in submerged culture (submerged h=12.0±1.3 μm; ALI h=22.0±1.8 μm; P<0.05), while the transepithelial electrical resistance (TEER) was not significantly affected by the ALI culture (**Fig 1B, C**). In immunostained Transwell inserts of Caco-2/BBe cells grown in submerged culture, DRA was primarily localized at the apical surface and under baseline conditions was generally distributed as small punctate structures (white arrowheads), while, in some areas, there were DRA aggregates (red arrowheads) (**Fig. 1D**). Exposure of confluent Caco-2/BBe cells to ALI for 2 days caused a major reorganization of DRA, which now was present in compact aggregates at the apical membrane. Multiple large aggregates of apical DRA are shown with the red arrowhead in the orthogonal view, XZ section, and in an XY section close to the top of the BB (**Fig ID**). ALI was associated with a threefold increase in DRA expression (3.5 ± 0.08-fold, both bands together) compared to cells grown in submerged cultures (F**ig. 1E**). The DRA higher molecular weight highly glycosylated band (ref from ming paper) was significantly increased with ALI 4.9± 0.5-fold) (**Fig. 1E**), while the core glycosylated band was not significantly altered (1.5 ± 2.2 fold) when compared with the cells grown submerged throughout. Importantly, the increase in DRA protein expression due to ALI modification was significantly greater than the change in DRA mRNA (**Fig. 1 F**). Caco-2/BBe DRA mRNA expression was increased 1.53 ± 0.3-fold in ALI culture compared to cells grown in submerged culture. Caco-2/BBe cells exposed to 4 and 6 days of ALI also had increased expression of DRA (**Supplementary Fig 1A, B**). However, Caco-2/BBe cells became multi-layered when exposed to more than 2 days of ALI culture (**Supplementary Fig 1B**). Therefore, the rest of the studies were performed on monolayers of Caco-2/BBe cells with 2 days of ALI.

**Figure 1.**
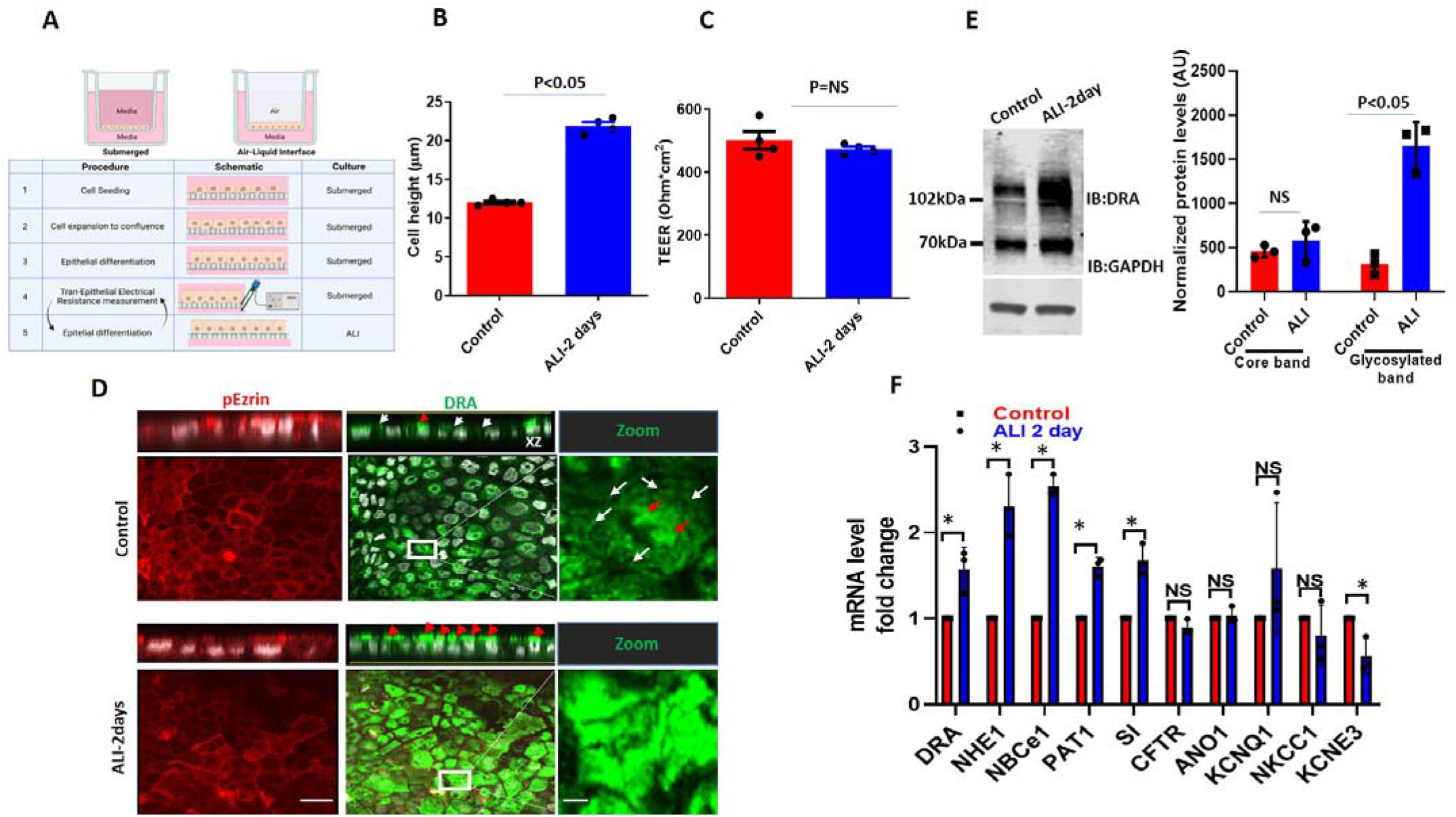
ALI culture increases DRA mRNA and protein expression in Caco-2/BBe cells. (**A**) A schematic overview of the procedures for creating ALI culture in intestinal epithelial cells including TEER measurement. (**B**) Cell height in Caco-2/BBe cells grown as a submerged culture for 12-14days (red) or followed by ALI-2days after 12 days post confluency (blue). (**C**) TEER of Caco-/2BBe monolayers from submerged and ALI culture. Mean±SEM of N=3-4 independent experiments is shown in **B** and **C**. P values represent unpaired Student’s t-test. (**D**) Confocal fluorescence microscopy of DRA (green), p-Ezrin (red), and nuclei (white). XZ orthogonal views (upper) and XY projections at the level of the apical membrane (lower). White arrowheads represent small aggregates of DRA-containing structures and red arrowheads represent large aggregates (platforms) of DRA on the apical side of Caco-2/BBe cells. Inserts show expanded areas. Scale bar: 10 μm and insert: 5 μm. (**E**) Left: IB of DRA and GAPDH of submerged (control) and ALI cultures. Right: densitometric analysis of DRA blots from N=3 independent experiments separately quantitating DRA upper and lower bands normalized to GAPDH. Results are Means ±SEM. (**F**) Quantitative real-time PCR analysis of changes in mRNA expression of multiple ion transport proteins and sucrase-isomaltase. Results are shown as Mean±SEM of fold change. N=3-5 independent experiment; P values represent unpaired Student’s t-test.

To investigate the alterations of other ion transport proteins after ALI, mRNA levels of selected ion transporters and the BB protein sucrase-isomaltase (SI) were also compared between submerged and ALI cultures. The ion transporters selected are known to play important roles in Cl^-^ and HCO_3_^-^ secretion, electroneutral Na^+^ absorption, and intracellular pH regulation. As shown in **Fig 1F** several ion transporters were significantly up-regulated at the mRNA level in Caco-2/BBe cells after 2-day ALI. These included *NHE1* (2.2-fold), electroneutral Na^+^/HCO_3_^-^ co-transporter 1 (*NBCe1*) (2.4-fold), and putative anion transporter 1 (*PAT-1*) (1.7-fold). Sucrase-isomaltase (*SI*), a brush-border glucosidase and intestinal differentiation marker, was also up-regulated significantly at the mRNA level in ALI cells. Some ion transporters were down-regulated significantly after ALI, including the potassium channel voltage-gated, subfamily E, regulatory subunit 3 (*KCNE3*) (2-fold). mRNA levels of the following ion transporters were not significantly changed by ALI: Anoctamin-1(ANO1), voltage-gated, subfamily Q, member 1 (*KCNQ1*), NKCC1, and CFTR (**Fig. 1F**). Overall, these results suggest that DRA resides at the apical side of enterocyte and that DRA undergoes extensive reorganization at the apical membrane along with an increase in expression during ALI culture, which includes both transcriptional changes as well as post-translational modifications. These changes are not specific to DRA, and ALI affects the expression of multiple apical and basolateral transport proteins and at least one BB digestive enzyme.

### ALI culture causes an increase in ASMase activity and colocalization with DRA

Aggregates like those of DRA in ALI-exposed Caco-2/BBe cells have been previously reported to be due to an increase in ceramide content in the plasma membrane^14^. Ceramide changes the biophysical properties of the membrane and causes lipid rafts to cluster together^15, 17^. ASMase is an enzyme that causes an increase in ceramide content in the plasma membrane by cleaving phosphorylcholine from sphingomyelin^18^ (**Fig 2A**). In many cells, ASMase is translocated from the lysosomal compartment to the cell surface in response to a broad range of pathological stimuli^19, 20^. The translocation of ASMase in response to stimuli was previously identified using immunostaining of ASMase under nonpermeabilized conditions^13^. To examine if ASMase is increased with ALI and if it is translocated to the BB during ALI culture, we compared ASMase immunostaining using confocal microscopy to examine the apical domain of unpermeabilized, well-differentiated Caco-2/BBe-Flag DRA cells grown submerged and after ALI exposure. ASMase immunofluorescence was readily detected in non-permeabilized Caco-2/BBe cells in both conditions, while increased immunofluorescence was detected after ALI stimulation. By immunostaining, Flag-DRA and ASMase co-localized under both submerged and ALI conditions. Moreover, increased localization of Flag-DRA and ASMase was observed at the apical domain (shown with white enclosed areas) after two days of ALI exposure to Caco-2/BBe cells (**Fig. 2B**). Manders overlap coefficient analysis showed 90% spatial colocalization between Flag-DRA and ASMase at the BB (**Fig. 2B**). The BB expression of DRA was compared between submerged and ALI-exposed Caco-2/BBe cells using cell surface biotinylation. The analysis of the surface-to-total ratio of DRA was significantly increased after ALI exposure as compared to submerged culture (**Fig 2C**). Since CFTR surface expression was previously reported to be elevated due to an increase in membrane ceramide levels in airway epithelial cells^14^, we also compared CFTR surface expression between submerged and ALI cultures in Caco-2/BBe cells. Similar to DRA, the surface expression of CFTR was significantly increased in Caco-2/BBe cells after 2-days of ALI exposure (**Fig 2C**). To determine if ASMase activity was stimulated by ALI, we used a fluorometric assay as described in Materials and Methods. ASMase activity was increased following ALI exposure (submerged: 1.5±0.5; ALI: 3.5±0.4; P<0.05 vs submerged) (**Fig. 2D**). These results indicate that ASMase is constitutively present on the exterior surface of the BB of Caco-2/BBe cells, and its amount and activity increase with ALI exposure as does an increase in DRA localization at the BB.

**Figure 2.**
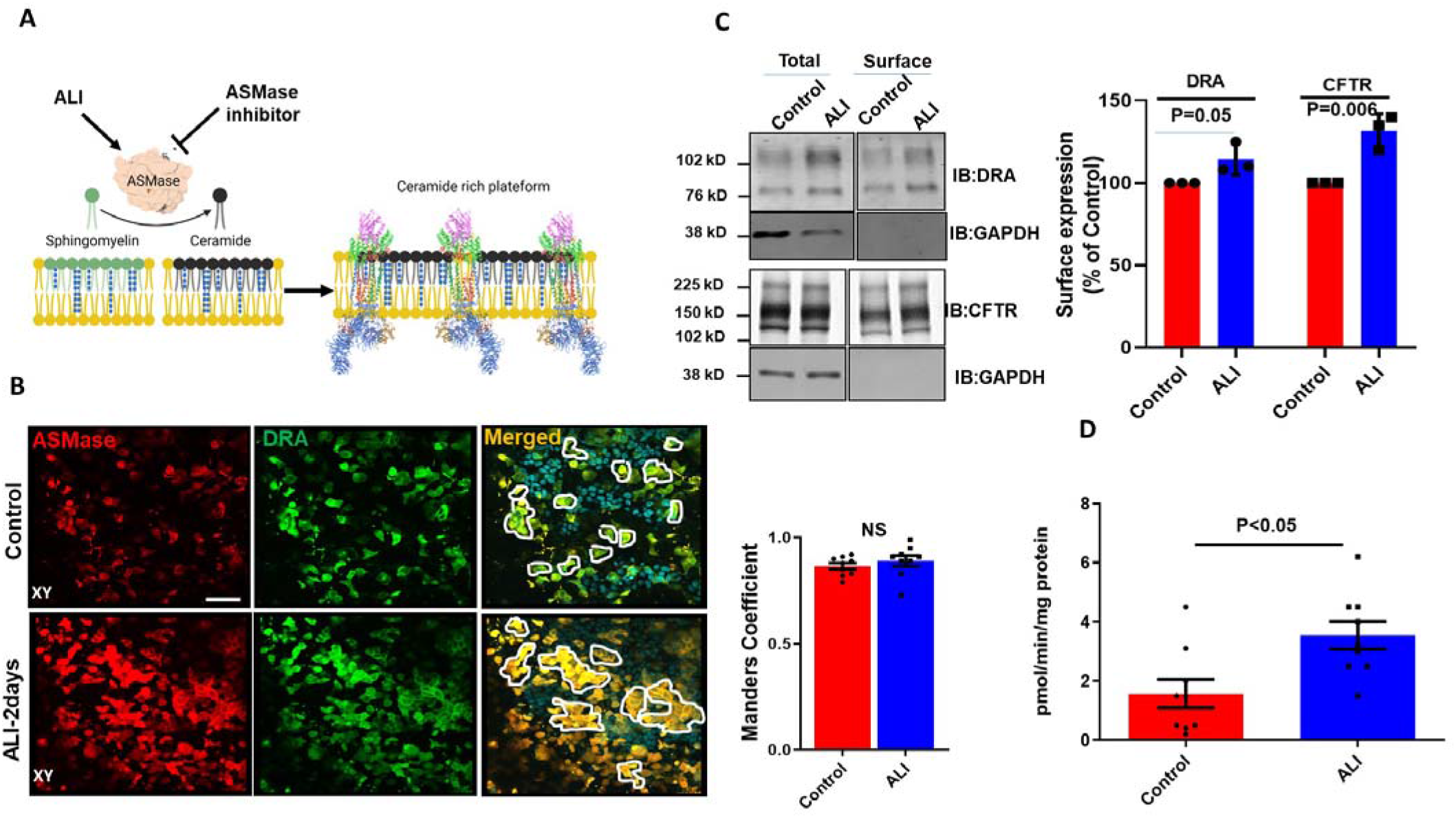
Increase in surface expression of DRA and ASMase activity in response to ALI. (**A**)Cartoon showing the role of ASMase in changing membrane ceramide amount and fusion of lipid rafts aggregates due to increase in membrane ceramide content. (**B**) Immunofluorescence detection of ASMase (red) and Flag-DRA (green) in the apical membrane of unpermeabilized, well-differentiated Caco-2/BBe cells grown as submerged cultures or followed by ALI-2days modification of 12-14days post confluency. Co-localization of endogenous ASMase and Flag-DRA was detected in cells grown as submerged cultures and was enhanced by ALI-2 days modification. A single plane (XY) at the apical surface of the Z-stack section is shown. Manders overlap coefficient showing 90% spatial colocalization of Flag-DRA and ASMase inside enclosed areas. Scale bar 50μm, N=3 repetition of the experiment shown were performed (**C**) Left: A representative immunoblot and densitometric analysis of total and cell surface biotinylation of DRA (above) and CFTR (below) in Caco-2/BBe cells grown as submerged or ALI-2days modification after12-14days post confluency. Right: quantitation of surface to total ratio normalized to GAPDH and expressed as a percent of control. Results are Mean ± SEM, N=3 independent experiments (**D**) Normalized ASMase activity in 12-14 days post confluent Caco-2/BBE cells and changes in activity after ALI-2 days modification. Results are Mean ± SEM, N=3-8 independent experiment. P values represent unpaired Student’s t-test.

### Basal and agonist-stimulated DRA activity in intestinal epithelial cells is increased by exposure to ALI culture

Basal DRA activity, measured by the initial rate of alkalinization that immediately follows the removal of Cl^-^ from the apical HCO_3_^-^ bathing solution, was compared between Caco-2/BBe cells grown as submerged culture or modified as ALI for 2-days. The initial rate of Cl ^-^ /HCO _3_^-^ exchange was significantly increased by ALI (135.0±1.5%) (**Fig. 3A, B**). We have recently shown that forskolin acutely stimulates DRA activity, which involves in part trafficking to the apical plasma membrane^21^. Therefore, we compared the response of forskolin between submerged and ALI-modified Caco-2/BBe cells. Forskolin (10 μmol/L, 10 min) significantly increased DRA activity in Caco-2/BBe submerged (127.0± 8.1%) and caused a non-significantly greater increase in ALI-modified cells (154.0± 0.5%). Previous studies in primary human bronchial epithelial cells suggested that secretagogues, including the peptide vasoactive intestinal peptide (VIP) and muscarinic agonist carbachol, enhance retention of transport protein (CFTR) on the plasma membrane by triggering membrane lipid conversion into ceramide and fusion of ceramide-rich platforms^13^. We, therefore, investigated if forskolin caused a fusion of DRA into clusters that looked similar to the demonstrated ceramide-rich platforms in human bronchial epithelial cells. Initially, the dimensions of the membrane microdomains that contain DRA were examined and the number of DRA clusters per unit area before and after forskolin stimulation was compared. Based on previous validations and nomenclature, membrane protein-containing areas having dimensions near the limit of optical resolution (<0.25 μm diameter) were considered clusters whereas large aggregates (≥2 μm dia.) were considered platforms_13, 14_. Quantitation in Caco-2/BBe cells showed that the cells grown in a submerged culture had higher numbers of DRA clusters (<0.25 μm) (average size:1.2±0.3 μm) compared to ALI-2days culture (average size: 4.8±0.3 μm) but a much smaller number of platforms (≥2 μm) (**Fig. 3C, D**). In submerged cultures, forskolin caused an increase in the number of platforms (average size 3.0±0.4 μm) and a significant decrease in the number of clusters. In contrast, in Caco-2/BBe cells after ALI exposure DRA was only present in platforms (≥2 μm dia) and the size of platforms further increased (average size: 6.1±0.4 μm) in response to forskolin (**Fig. 3D).** We also compared the Fsk-induced short-circuit increase between submerged and ALI cultures in Caco-2/BBe cells. Similar to reports in primary human bronchial epithelial cells, there was a significant increase in forskolin-stimulated CFTR activity (**Fig. 3E**) after 2days of ALI exposure. These results suggest that an increase in the size of DRA-containing clusters at the plasma membrane to form platforms due to ALI may contribute to the increase in basal and forskolin-stimulated DRA activity in these cells.

**Figure 3.**
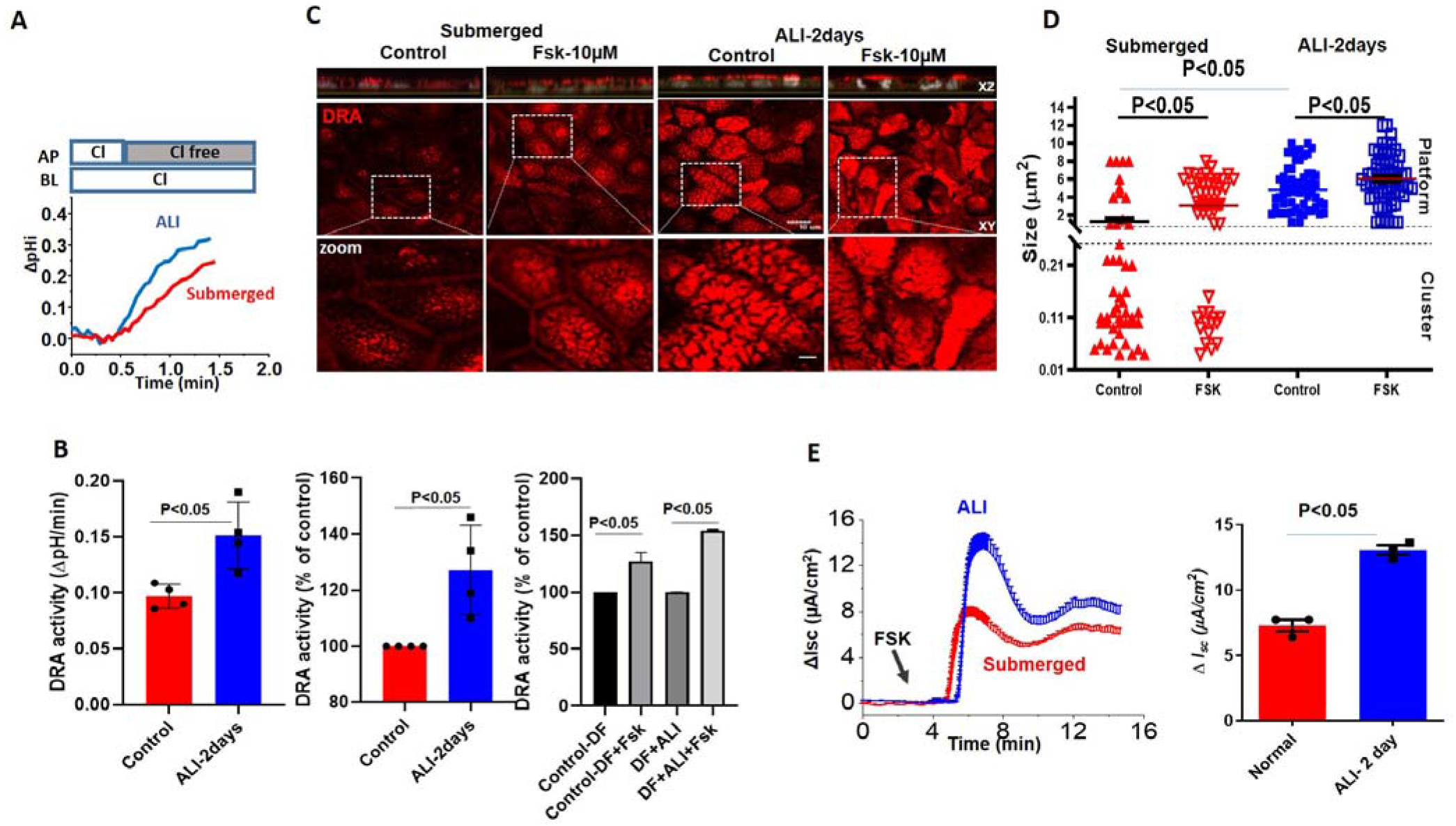
Increase in DRA basal activity and agonist stimulation in Caco-2/BBe cells after ALI exposure. Difference in basal and stimulated DRA activity and expression between Caco-2BBe cells grown as submerged culture for 12-14days (red) or followed by ALI-2days (blue) after 12 days post confluency: (**A**)A representative trace and (**B**)Basal Cl^-^/HCO_3_^-^ exchange activity and Forskolin stimulation. Left initial rates as ΔpH/min, center and right normalized as percent of control, Results are Mean ± SEM, n=3 independent experiments. P values represent unpaired Student’s t-test. (**C**) Left: A representative confocal image showing changes in sizes of endogenous DRA-containing clusters and appearance of the platform (clusters>2μm) comparing submerged and ALI exposure culture before and after forskolin (10 μM, 10 min) stimulation. All specimens were immunostained for DRA (red) and nuclei (grey) with XZ sections (above), XY sections at the level of BB (middle), and zoomed view of the boxed area below. Scale bars: 10 μm, insert 5 μm. (**D**) Quantitation of DRA-containing areas as in (C). DRA-containing areas having dimensions near the limit of optical resolution (<0.25 μm) were considered clusters whereas large aggregates (≥2 μm dia.) were considered platforms. Size of aggregates in μm calculated for ~100 individual clusters and platforms from three areas/per monolayer/condition. Results are Mean ± SEM, n=3-8 independent experiment. P values represent unpaired Student’s t-test. (**E**) A representative trace (left), and summary (right) of ΔI_sc_ response showing forskolin-induced Δi_sc_ from submerged culture and ALI-2days exposed monolayers after 12-14days confluency. Results are Mean ± SEM, *n* = 8–12 filters/condition from 3 independent experiments. P values represent unpaired Student’s t-test.

### ASMase inhibitor Desipramine treatment reduces ASMase and basal DRA expression as well as ALI-induced increase in DRA amount at the BB

DRA is in part localized in the lipid raft pools of the Caco-2/BBe plasma membrane^7, 22^. Since ASMase and DRA co-localize and are present in increased amounts in apical membrane platforms in ALI, the dependence of DRA abundance in the plasma membrane on ASMase was determined. Desipramine, which functionally inhibits ASMase, was used^23^. Caco-2/BBe cells were transduced with adenoviral Flag-DRA, then 24h later the cells were exposed to desipramine for 1 hr (13μM), followed by exposure to ALI for 2 days in the continued presence of desipramine (13μM). As shown in **Fig 4A and B**, Desipramine decreased both basal ASMase amount and activity and the increases caused by ALI. Desipramine pretreatment also decreased the basal expression as well as ALI-induced increase in BB amount of Flag-DRA (**Fig. 4A**). Importantly, pretreating cells with desipramine for 1 hr reduced ASMase activity below the control level, suggesting that there is ASMase activity in Caco-2/BBe cells under basal conditions. These results indicate that ALI-induced accumulation of DRA at the apical membrane depends on the activity of ASMase.

**Figure 4:**
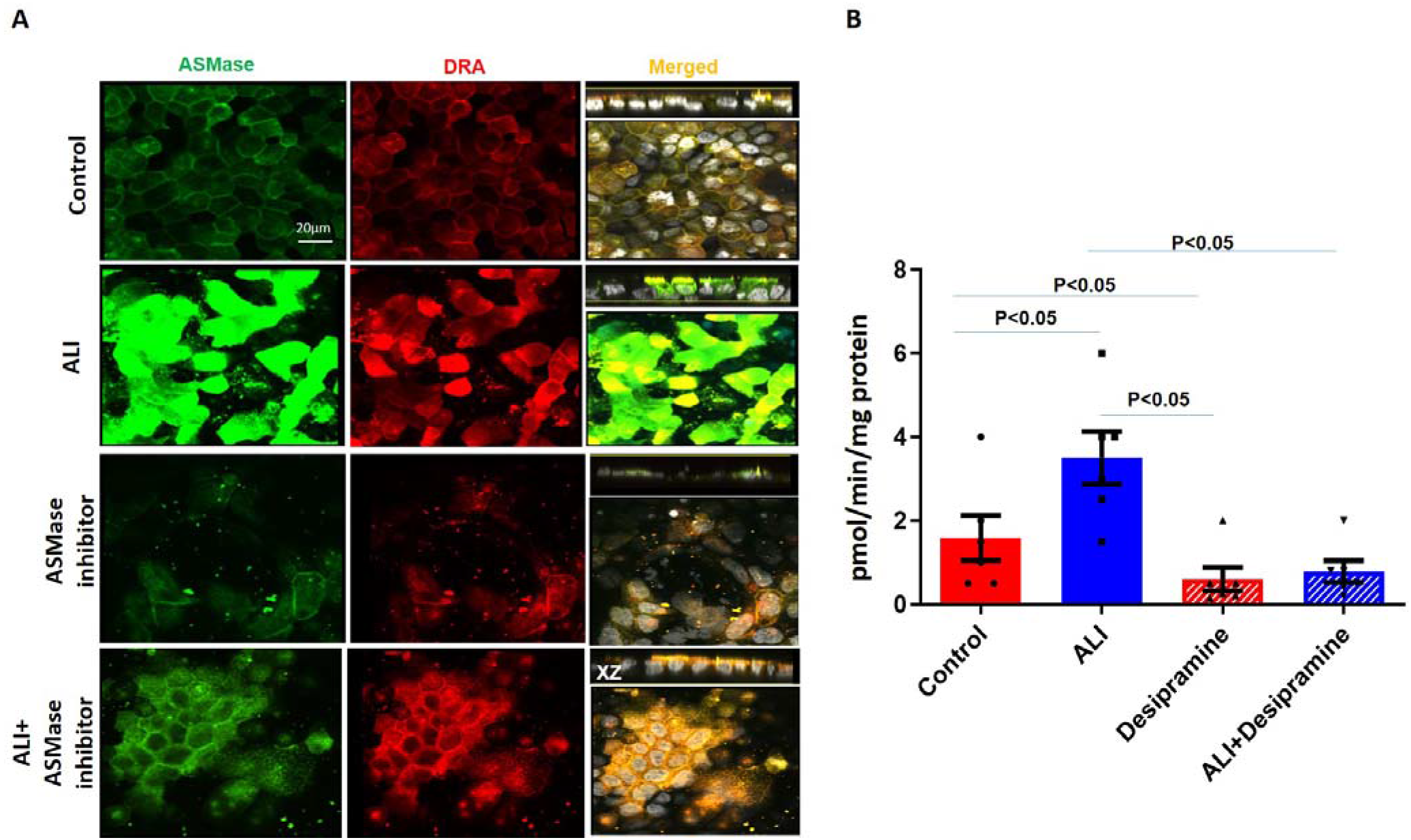
Increase in surface DRA expression due to ALI was blocked by the ASMase inhibitor, Desipramine. The effect of the ASMase inhibitor, Desipramine was determined on apical plasma membrane localization of DRA in Caco-2/BBe cells grown as a submerged culture for 12 days or followed by ALI-2 days. Flag-DRA was transduced into Caco-2/BBe cells, 24h before ALI initiation. Transduced cells were exposed to ALI-2 days or were pretreated with 13μM **Desipramine 1hr** before ALI was initiated and during the ALI. (**A**) A representative confocal image after immunostaining for Flag-DRA (red), ASMase (green), and nuclei (white). A single plane of multi Z-stack is shown. All panels represent XY projection near the top of the cell, the right panel includes XZ projection on the top. Scale bar 20 μm. (**B**) Normalized ASMase activity in Caco-2/BBe cells expressing Flag-DRA is shown for four conditions in A. Pretreating cells with 13μM **Desipramine** reduced ASMase activity below the control level, suggesting there is some ASMase activity under basal conditions. Results are Mean ± SEM; N=3-8 independent experiments. Statistical analysis was performed using ANOVA.

### ALI causes an increase in the detergent-insoluble fraction (lipid raft) of DRA in intestinal epithelial cells

Lipid rafts are enriched with cholesterol and sphingolipids and are resistant to detergent solubilization such as by Triton X-100; hence they are included in the detergent-insoluble (DI) membrane fraction^7^. Since insolubility depends on the cholesterol composition of these microdomains, we determined the effect of methyl-β-cyclodextrin (MβCD) (selectively removes most of the cholesterol from the membrane) on the association of DRA with the DI fraction). Total membrane from DMSO or MβCD (10μM) treated submerged Caco-2/BBE cells were incubated with Triton-X 100 at 4°C and the DI and DS fractions were separated and analyzed using western blot analysis. As shown in Fig. 5A, the DI fraction of the total membrane prepared from Caco-2/BBe cells expressed more DRA compared to the DS fraction under basal conditions. Moreover, the treatment with MβCD caused a significant reduction in the expression of DRA in the DI membrane fraction (**Fig 5A**). These findings support previous reports of the presence of DRA in lipid rafts in the total membrane of Caco-2/BBe cells^7, 22^. Whether ALI altered the association of DRA with lipid rafts or the DI fraction of Caco-2/BBe total membrane was determined. As shown in Fig. **5B**, a higher amount of DRA was expressed in the DI compared to the DS fraction of both the submerged (called control) and ALI-exposed Caco-2/BBe cells. However, DRA expression in the DI fraction was significantly enhanced in the ALI compared with the submerged culture. These findings suggest that ALI increases the amount of DRA in the lipid raft fraction of the plasma membrane.

**Figure 5:**
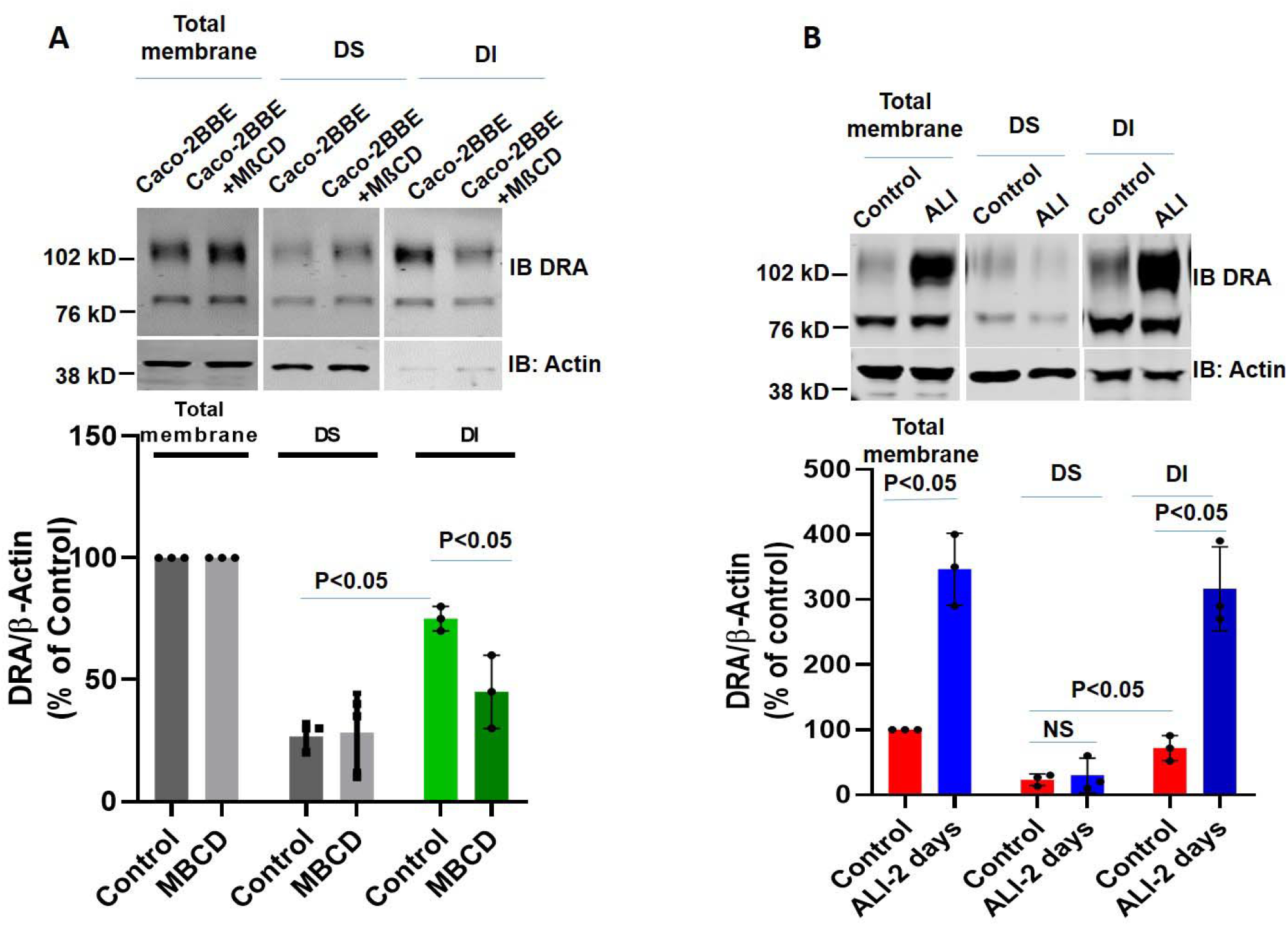
ALI causes an increase in the detergent-insoluble fraction of DRA in intestinal epithelial cells. DRA is associated primarily with the detergent-insoluble (DI) fractions of Caco-2/BBe cells and ALI-2 days exposure causes an increase in the DI fraction of DRA of total membranes. (**A**) DI vs DS distribution of DRA in the presence or absence of methyl-β-cyclodextrin (MβCD; 10 μM, 1 h). The single experiment above and Mean ±SEM of N=3 similar experiments are shown below. (**B**) Total membranes from submerged and ALI-2days exposed cells, were solubilized in a buffer containing Triton X-100 and then detergent-soluble (DS) and DI fractions were isolated as described in MATERIALS AND METHODS. Equal amounts of proteins (~50 μg) from DI and DS fractions were separated on 10% PAGE and then analyzed by Western blotting for DRA and actin expression. The data were quantified by densitometric analysis and expressed as a percent of control and fold change representing Means ± SEM of 4 determinations. P values represent unpaired Student’s t-test.

### ALI enhances the differentiation of human colonoids

Caco-2/BBe cells are human colon cancer-derived cell lines. To determine whether ALI also affects DRA in normal human intestinal epithelial cells, we used colon-derived normal human intestinal enteroids also called colonoids (HIEs). When HIEs are differentiated, they include the different epithelial cell types present in the normal intestine, and studies were done in monolayers of differentiated HIEs because DRA is expressed almost entirely in this population^21, 24, 25^. The monolayers were initially maintained in the undifferentiated (UD) crypt-like state by growth in Wnt3A, Rspo-1, and Noggin, and for these studies were caused to differentiate (DF) by the withdrawal of growth factors (Wnt3A and Rspo-1) for 5-6 days^24, 26^. DRA expression was compared under 4 different conditions: 1) UD, 2) UD+ALI-5days, 3) DF, and 4) DF+ALI-5days. UD enteroid monolayers from submerged or ALI conditions did not exhibit a significant time dependent increase in TEER. In contrast, the TEER of the differentiating monolayers from both the submerged and the ALI conditions increased significantly and similarly over time (**Fig 6A**). In these studies, even after 5days of ALI exposure, colonoid monolayers did not form multilayers, as seen in Caco-2/BBe cells (supplementary **fig 1B).**

**Figure 6:**
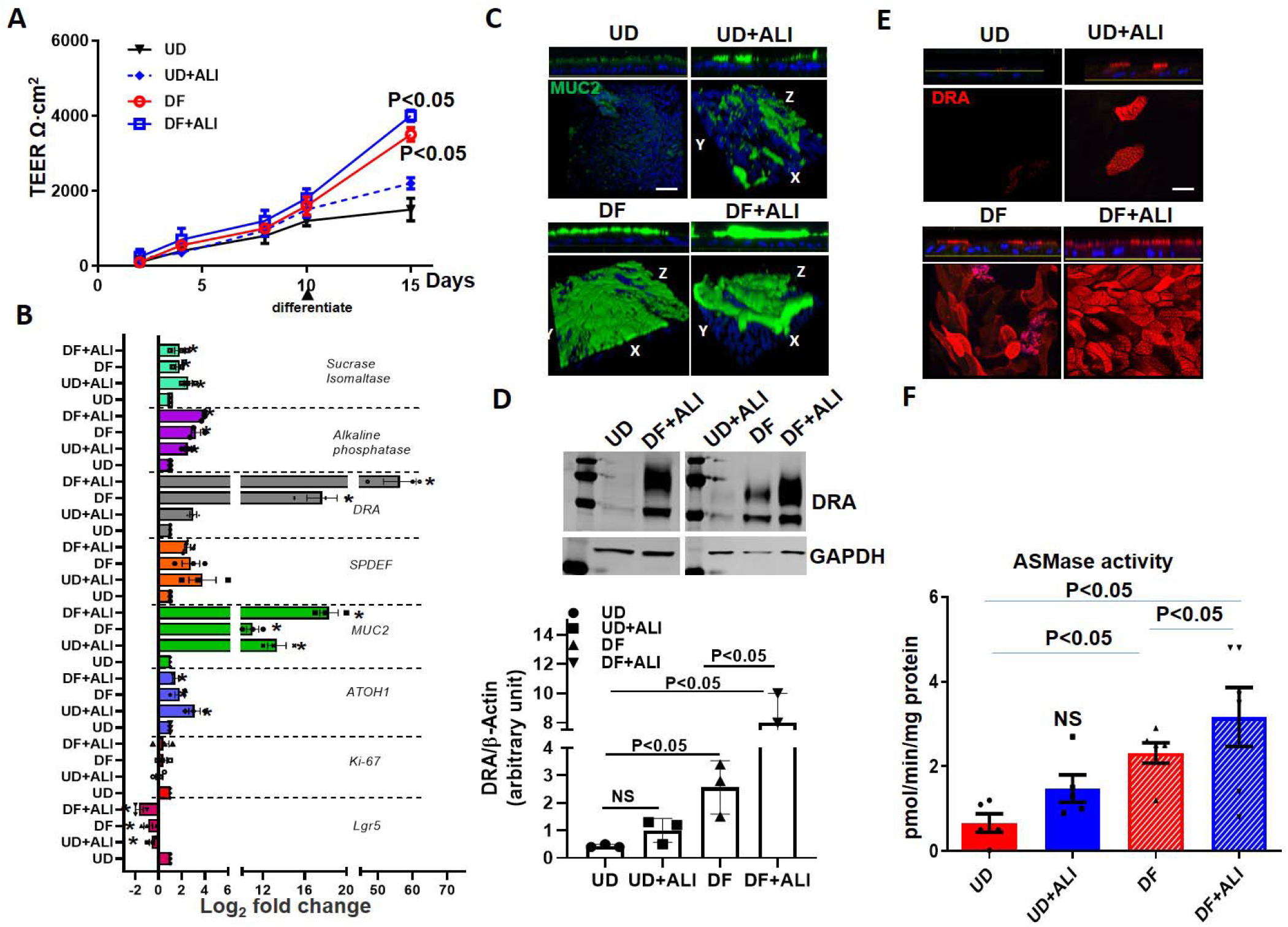
ALI Increased differentiation of human colonoids. Human intestinal colonic organoids (colonoids) derived from biopsies were grown as confluent epithelial monolayers on permeable inserts. Post confluency monolayers studied in all these experiments compared: 1) UD; 2) UD + ALI; 3) 5 days DF; 4) 5 days DF + ALI. (**A**) Changes in TEER in response to differentiation or ALI-modification are shown. Results are Means ±SEM of N=3-4 experiments. P value is compared with the UD control. (**B**) Relative mRNA levels of genes used to evaluate proliferation and differentiation including DRA by qRT-PCR. Messenger RNA levels are normalized to 18S ribosomal RNA expression. Results are normalized to UD set as 1 and expressed as Log_2_ fold change. Data were analyzed using a two-tailed Student’s t-test with Welch’s correction. *\*P* < 0.05 compared with UD control. Results are Means ±SEM of N=3-4 independent experiments. Atoh, atonal homolog 1; Lgr5, leucine-rich repeat-containing G protein-coupled receptor; Muc2, mucin 2; ki67, nuclear protein ki67; SPDEF, SAM Pointed Domain Containing ETS Transcription Factor. (**C**) Methanol–Carnoy’s fixed colonoid monolayers stained with anti-MUC2 (*green*), and nucleus stained with DAPI (*blue*). Representative confocal XZ (*above*) and 3D-XYZ (*below*) projections depicting the MUC2 layer in colonoid monolayers is shown. Scale bar 10 μm. (**D**) A representative Western blot and densitometry analysis from multiple experiments shows changes in DRA protein expression. Results are Means ±SEM of N=3 experiments. (**E**) Confocal fluorescence microscopy of DRA (red) expression in colonoids. Upper, XZ projection and lower, XY projection at the level of apical domain. Scale bar 10 μm. (**F**) Normalized ASMase activity in colonoids. Results are means ± SEM of 3-5 independent experiments. P value is compared with the UD control. P values represent unpaired Student’s t-test.

To investigate if ALI altered the differentiation of the colonoids, quantitative RT-PCR expression of a selected panel of genes was performed on monolayers collected from the above 4 conditions. As shown in **Fig 6B**. Lgr5 and Ki67 mRNA were reduced in all conditions except the UD enteroids. Similarly, atonal homolog 1 (ATOH1) and mucin 2 (MUC2) expression were increased in all conditions compared with UD monolayers. Additional markers of enterocyte differentiation were examined. Sucrase-isomaltase and alkaline phosphatase increased similarly in all conditions except in the UD monolayers^27^. Further support for the effect of ALI on enteroid differentiation is demonstrated in **Fig 6C** in which the mucus layer of the enteroids was determined by confocal microscopy. ALI increased the thickness of the mucus layer in the UD + ALI condition and DF+ALI monolayers with the largest mucus layer present in the DF + ALI condition.

DRA mRNA expression was significantly increased in UD+ALI-5days (~2.5fold vs UD), and DF (~15folds vs UD) monolayers. However, cells from DF+ALI-5days monolayers showed a very large upregulation of DRA mRNA expression (~60folds vs UD). The DRA protein expression was also measured using western blot analysis of lysates prepared from the colonoids (**Fig 6D**). The DRA protein expression tracked the changes in mRNA being significantly increased by DF and DF + ALI conditions, with the largest change in the DF + ALI condition, an increase compared to UD of 18 fold while DRA expression in the DF condition was increased by 5-fold vs UD. UD + ALI caused a slight but not significant increase in DRA protein expression. In addition, the apical membrane expression of DRA was increased in all three conditions compared to the UD enteroids with the greatest increase being in the DF + ALI conditions (**Fig 6E**). The ASMase activity in cells under the above 4 conditions was also determined (**Fig 6F**). Compared to UD monolayers, which had very low ASMase activity, monolayers from the other 3 conditions had higher ASMase activity, but these were only statistically significant for the DF and DF + ALI conditions, with the highest ASMase activity in the DF+ ALI condition (**Fig 6F**). Overall, these results suggest that exposure of monolayers to ALI enhances the differentiation of colonoids which is associated with an increase in DRA and ASMase expression. The increase in differentiation occurred much more in monolayers in which differentiation was initiated by growth factor removal and the changes were much less in undifferentiated enteroids.

### DRA expression and activity are further enhanced by ALI culture of differentiated human colonoids

To explore the role of ALI in enhancing the differentiation of already differentiated human colonoids, we exposed enteroids differentiated for 4 days to 2 subsequent days of ALI and studied changes in specific gene expression including DRA. These studies were performed in the continued presence of DF media on the basolateral side throughout. Changes in DRA expression were compared between colonoids from 4 days DF+ALI-2 days with 6 days DF. The TEER was not significantly different between 6-day DF and 4dayDF+2dayALI culture (**Fig 7A**). In contrast, the mRNA expression of differentiation markers including sucrase-isomaltase and alkaline phosphatase further increased along with DRA which had a small but significant increase in the ALI condition (~1.4folds vs 6dayDF) as compared to 6dayDF (**Fig 7B**). Of note, the protein expression of DRA was ~3folds higher in 4dayDF+2dayALI culture as compared to 6dayDF culture (**Fig 7C**). Accordingly, basal DRA activity was ~1.6folds higher in 4dayDF+2dayALI culture (**Fig. 7D**). We also compared the stimulation of forskolin on monolayers under these 2 conditions. The percentage of stimulation of DRA in response to forskolin was not significantly different between the 4dayDF+2dayALI culture and 6dayDF culture (DF:150± 0.3%; DF+ALI: 137± 0.1%; P=n.s vs DF-6days) (**Fig 7E**). Immunofluorescence analysis of monolayers also showed higher expression of DRA on the apical membrane and larger DRA-containing aggregates in 4dayDF+2dayALI culture (**Fig 7F**). Moreover, significantly higher ASMase activity was detected in cells from 4dayDF+2dayALI culture as compared to DF-6 days and both were significantly higher than ASMase activity in UD monolayers (**Fig 7G**). These results show that DF of colonoids increases ASMase activity as well as DRA expression and that exposure of DF colonoids to ALI further increases ASMase activity along with DRA expression.

**Figure 7:**
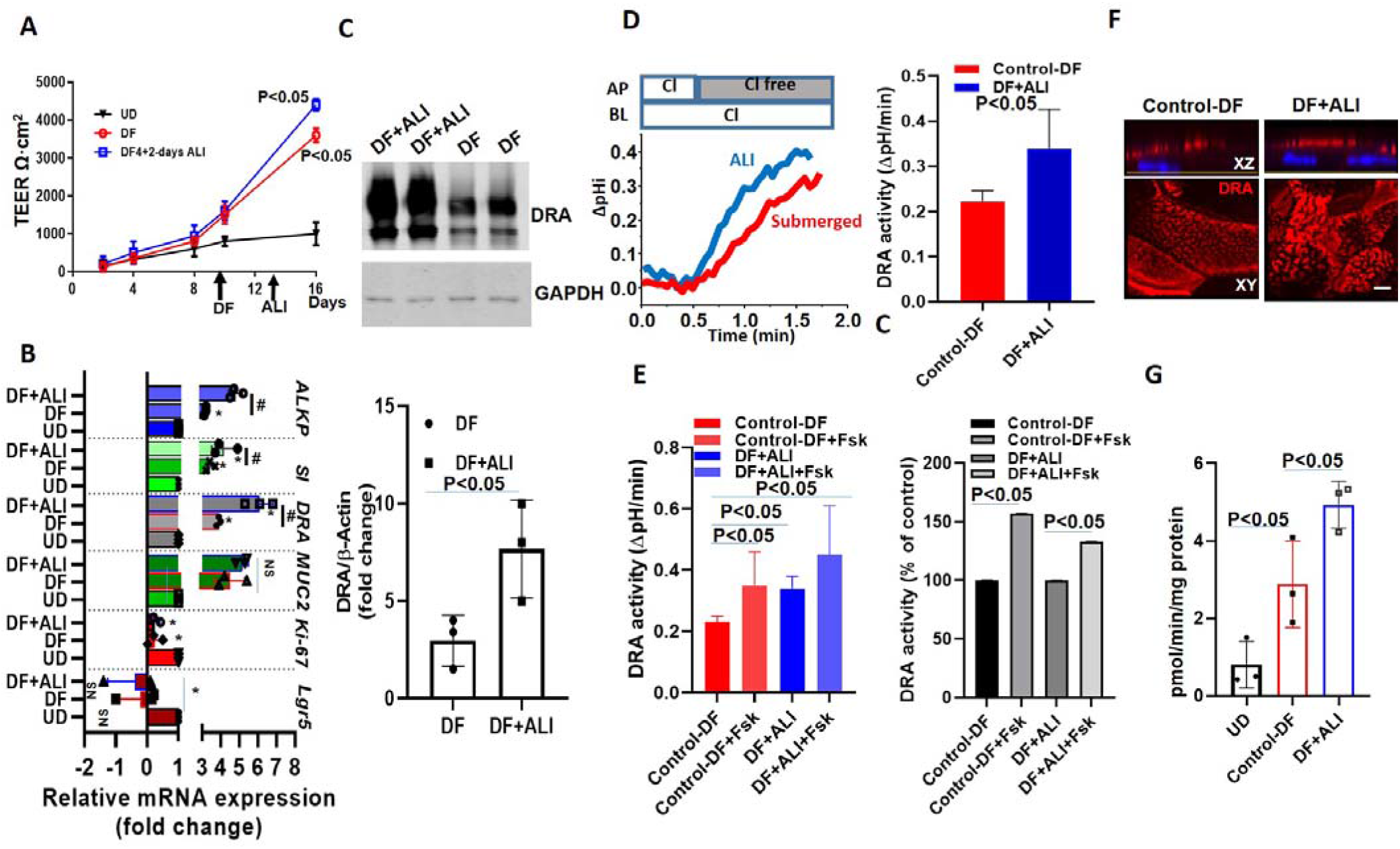
DRA expression and activity were further enhanced by exposing differentiated human colonoids to ALI culture. Human colonoids were grown as confluent epithelial monolayers on permeable inserts. Post confluency monolayers were exposed to either differentiation medium (DF) for 6 days or to 4-days DF followed by 2-days ALI-modification. (**A**) Changes in TEER in response to differentiation or ALI-modification are shown. Points are Means ± SEM, N=4. Relative mRNA expression of DRA by quantitative PCR. Messenger RNA levels were normalized to 18S ribosomal RNA expression. The results are normalized to UD set as 1 and expressed as fold change. Results are Mean ± SEM from N=3 independent experiments. *P<0.05 vs UD control, ^#^P<0.05 vs 6days DF control. P values were analyzed using two-tailed Student’s t-test with Welch’s correction. (**C**)A representative Western blot (above) and densitometry analysis Means ± SEM (below) from multiple experiments (N=3) showing an increase in DRA protein expression in 4-days DF+2-days ALI compared to 6 days DF condition. P values represent unpaired Student’s t-test. (**D**) A representative trace of basal Cl^-^/HCO_3_^-^ exchange activity (left) and Means ± SEM of N=3 independent experiments (right) and (**E**) Basal DRA activity and forskolin (10μM, 10mins) stimulation comparing 6 days DF and 4-days DF+2-days ALI conditions, shown as DRA activity and as a percent of control, N=3. Statistical analysis was performed using ANOVA. (**F**) Confocal fluorescence microscopy of DRA (red) and nucleus stained with DAPI (blue) expression in colonoids; upper XZ and lower XY project at the level of apical domain. Scale bar 10 μm. (**G**) Normalized ASMase activity in colonoids comparing UD, 6-day DF, and 4 day DF + 2day ALI. Results are Means ± SEM, N= 3 independent experiments. P values represent unpaired Student’s t-test.

## Discussion

In this study, we showed that ALI exposure enhances differentiation of human proximal colonic enteroids and Caco-2 cells; this occurred in previously differentiated enteroids as well as when cells were exposed to ALI during growth factor removal-induced differentiation and, although to a lesser extent, in undifferentiated enteroids. The aspects of differentiation that were affected included reduced expression of stem cell markers, including mRNA expression of *LGR5*, reduced proliferation (*Ki67*), increased expression of *ATOH1*, along with the goblet cell marker *MUC2*, and a thicker apical mucus layer. Additionally, there was increased expression of apical markers in villus cells (sucrase-isomaltase, and alkaline phosphatase), as well as DRA, which is normally present only in enterocytes that are at least partially differentiated and greatly increases in amount with differentiation. Similar changes also occurred in UD enteroids when exposed to ALI, although the extent of the increase was less and often failed to reach statistical significance. These changes included significantly reduced mRNA for *LGR5*, and increased *ATOH1*, and *MUC2* expression along with a thicker colonic mucus layer. Our previous findings in human duodenal enteroids suggested that differentiated (villus-like) enteroids express more DRA, NBCe1, and sucrase-isomaltase and reduced KCNE3 and NKCC1 expression^25^, all changes are similar to the effects caused by ALI in the current study. Together, these findings show that ALI enhances the differentiation of human colonoids. ALI is a standard method for the differentiation of human bronchial epithelial cells and it was recently shown that ALI-like effects can be achieved in submerged culture by controlling the oxygen content^28, 29^. The latter findings suggest that oxygen levels play a significant role in the differentiation of epithelial cells. While the effects of oxygen levels on enteroids differentiation have not been directly studied, our current findings are supported by previous studies suggesting that long-term culturing of organoids as submerged monolayers generated low oxygen tension, leading to cellular stress, reduced differentiation, and a lack of secretory cells. Notably, these effects were reversed by ALI culture which increases the oxygen levels of the enteroid monolayers and re-established healthy and differentiated monolayers^30–32^.

In this study, we emphasized two proteins that underwent increased expression as part of ALI-induced differentiation: ASMase and DRA. ASMase is an enzyme that breaks sphingomyelin into ceramide and phosphorylcholine in the outer leaflet of the plasma membrane. Ceramide changes the biophysical properties of membranes and causes lipid rafts to cluster together to form ceramide-rich platforms (CRP) previously described in airway epithelial cells, lymphocytes, and fibroblasts^14, 19^. The *in vitro* studies revealed that the generation of ceramide is sufficient to trigger the formation of distinct platforms even in purely artificial membranes without any cytoskeleton or other cellular proteins^33^. These platforms selectively trap or exclude specific proteins for biophysical and energetic reasons, and thus serve as a sorting unit for receptors and signaling molecules^19^. ALI increased ASMase activity in human colonoids and Caco-2/BBE cells and visibly enlarged apical membrane ASMase containing aggregates that fit the description of ceramide-rich-platforms^14, 34^. ASMase is ubiquitously expressed and localizes preferentially to the endo-lysosomal compartment. However, under certain conditions, it translocates to the plasma membrane^35^. In ALI of Caco-2/BBe cells, the plasma membrane translocation that occurred was demonstrated based on the ability to see its presence at the apical surface in non-permeabilized cells. Under basal conditions, ASMase co-localized with DRA in apical membrane clusters, and with ALI these clusters aggregated into CRPs that contained DRA. How ALI-induced activation of ASMase models normal physiologic or pathologic regulation of ASMase has not been established. ASMase is considered to have a protective role during infection based on the studies of ASMase knockout mice, which had exaggerated IL-1β release and increased mortality when infected with *Pseudomonas aeruginosa^36^.* ASMase is activated in response to various stress stimuli including 1) ligation of death receptors (tumor necrosis factor-α, CD95, and TRAIL), 2) radiation (UV-C and ionizing radiation), 3) chemotherapeutic agents (cisplatin, doxorubicin, paclitaxel, and histone deacetylase inhibitors), 4) viral, bacterial, and parasitic pathogens (rhinoviruses, Neisseria gonorrhea, Staphylococcus aureus, Pseudomonas aeruginosa, and Cryptosporidium parvum), and 5) cytokines (e.g. IL-1β)^20, 37–40^. Evidence from these studies in cell culture and animal models has been interpreted to suggest that activation of ASMase takes part in regulating cellular differentiation (monocyte to macrophages), growth arrest, apoptosis, and immune defense mechanisms^41, 42^. However, its possible role in stimulating enterocyte differentiation has not been suggested. Changes in enterocyte differentiation occur in multiple models of intestinal pathology, including infection by bacteria, viruses, and worms as well conditions of chronic inflammation. Given that DRA activity has been altered in several of these models, it is worth determining whether ASMase activation/plasma membrane translocation has a pathophysiologic role; and given its presence under basal conditions in the Caco-2/BBe plasma membrane, its role in regulating basal DRA activity under normal physiologic conditions that include the post-prandial state will be important to understand as well.

In parallel with the changes in ASMase, 2 days of ALI significantly increased the amount of total and apical membrane DRA and increased the localization of DRA in the detergent-insoluble portion of the total membrane and apical membrane. The increased DRA was almost entirely of the upper heavily glycosylated band with minimal change in the core glycosylated band, consistent with effects primarily on the apical membrane DRA pool^21^. These results are consistent with previous reports in Caco-2/BBe cells that showed that the majority of DRA is present in the detergent-insoluble membrane fraction^7^. An increase in DRA expression in human colonoids after a 6-day ALI culture was primarily due to transcriptional upregulation of the DRA gene expression due to enhanced differentiation of colonoids. However, when 4-day DF colonoids were exposed to 2-days ALI they showed enhanced DRA protein expression and activity, with less transcriptional upregulation. This effect was similar to Caco-2/BBe cells where ALI showed a higher effect on translational upregulation than via transcriptional stimulation (**Fig 1C**). In airway cells, enhancement of CRPs during regulation by physiological agonists is known to increase CFTR activity^13^. Similarly, in Caco2-BBe cells we found a significant increase in CFTR apical expression (**Fig. 2C**) and activity after 2days of ALI exposure (**Fig. 3E**). This suggests that there is a common mechanism of activation of CFTR by these platforms in both the pulmonary and intestinal cells as well as a similar mechanism by which ALI affects DRA and CFTR in human enteroids. We recently reported that forskolin stimulation of DRA is at least in part dependent on CFTR protein expression, supporting the studies of Ko and Muallem in oocytes^21, 43^. It remains to be determined if the CRPs have a role in the stimulation of DRA activity in response to forskolin in addition to the potential increase in the accessibility that results from having both DRA and CFTR present in the CRPs. Further studies are required to understand the regulation of DRA under physiologic and pathologic conditions that are associated with increased amounts of CRPs.

In summary, the present results with ALI demonstrated y that it causes enterocyte differentiation. Concerning the relevance of these findings to normal physiology, the intestine is primarily exposed to large luminal volumes related to eating, while much less luminal fluid is present during many hours of fasting or sleep. In this study, we have identified a new pathway for DRA stimulation that extends our understanding of apical compartmentalized DRA and potentially provides insights into how it interacts with CFTR. The physiologic and pathophysiologic significance and factors involved in the increase in ASMase activity in the plasma membrane remain to be established, although these results strongly support a role for ASMase activation in DRA and CFTR regulation. Moreover, understanding the distribution and lateral mobility of DRA in ceramide-rich platforms in the plasma membrane and how these aspects of DRA function are regulated by secretagogues will provide insights into understanding the role of the CRPs in the regulation of multiple brush border ion transport processes.

## Materials and methods

### Chemicals and reagents

were purchased from Thermo Fisher (Waltham, MA) or Sigma-Aldrich (St. Louis, MO) unless otherwise specified.

### Cell Lines

Caco-2/BBe cells were cultured in Dulbecco’s modified Eagle medium supplemented with 25 mmol/L NaHCO _3_, 0.1 mmol/L nonessential amino acids, 10% fetal bovine serum, 4 mmol/L glutamine, 100 U/mL penicillin, and 100 μg/mL streptomycin in a 5% CO _2_ /95% air atmosphere at 37°C. For experiments, cells were plated on Transwell inserts (Corning, Inc, Corning, NY) and studied 14-18 days after reaching confluency. The plasmid p3xFLAG-DRA was cloned into the adenoviral shuttle vector ADLOX.HTM under the control of a cytomegalovirus (CMV) promoter^25^.

Endoscopic specimens of the human proximal colon from healthy human subjects undergoing endoscopies for medically indicated conditions were used to establish primary cultures of the human colon, called colonoids, as previously described^44^. Colonoids were expanded and plated on Transwell inserts (polyester membrane with 0.4-μm pores Corning) to form monolayers, as previously described^21, 25, 26, 44^. The formation of colonoid monolayers was monitored by the measurement of TEER. Monolayers were maintained in an undifferentiated state by exposure to WNT3A, Rspon-1 and Noggin^26^. For differentiation, colonoids were maintained in a medium that lacked WNT3A, R-spondin1, and SB202190 for 5 days. Five days later, paired UD and DF enteroid monolayers were studied. Most results of the current study were obtained from colonoids derived from 1 healthy donor, with similar results observed in colonoids from 2 other donors. The procurement and study of human colonoids were approved by the Institutional Review Board of Johns Hopkins University School of Medicine (NA_00038329).

For air-liquid interface studies, apical culture media were removed for specified days from 12-14 days post confluent Caco2/BBe cells or confluent colonoid monolayers.

#### Immunofluorescence

Cells were grown on collagen-coated Transwell supports as submerged culture or exposed to an air-liquid interface for a specified number of days. To visualize DRA and the effects of ALI on DRA, cells were washed with PBS and fixed in 4% paraformaldehyde for 30-45 minutes, incubated with 5% bovine serum albumin/0.1% saponin in phosphate-buffered saline for 1 hour, and incubated with primary antibody against Flag or DRA (mouse monoclonal, 1:100, sc-376187; Santa Cruz, Dallas, TX) alone or in combination with a rabbit anti-ASMase antibody (1:200; Abcam ab227966) overnight at 4°C. Cells were then exposed to Alexa Fluor 488 goat anti-mouse and Alexa Fluor 594 goat anti-rabbit secondary antibodies (1:1,000 dilution; Invitrogen) for 1 h and mounted in ProLong Diamond Antifade Mountant (Invitrogen). Images were collected using ×40 oil immersion objective on an FV3000 confocal microscope (Olympus, Tokyo, Japan) with Olympus software and ImageJ software (NIH). For quantitative analysis, the same settings were used to image all samples. To examine ASMase and DRA distribution on the apical surface of Caco-2/BBE cells, the above procedure was followed without the membrane permeabilization step. The effect of ASMase inhibitor desipramine on ASMase and DRA distribution was assessed by pretreating cells on both sides of Transwells for 1 h and then exposing them to 2days-ALI in presence of desipramine. Individual clusters (optical resolution (<0.25 μm)) and platforms (large aggregates (≥2 μm dia.)) were encircled using ImageJ software, and their areas were estimated from the number of pixels × single pixel area^13, 14^. Cluster size was quantified using pixel number and total area. Microdomain areas were estimated by counting three areas/monolayers/conditions from three different experiments.

Analysis of MUC2 by immunofluorescence and confocal microscopy was carried out as previously reported^44–46^. Briefly, human colonoid monolayers were fixed with Carnoy’s solution (90% [v/v] methanol, 10% [v/v] glacial acetic acid), washed 3 times with phosphate-buffered saline, permeabilized with 0.1% saponin, and blocked with 2% bovine serum albumin + 15% fetal bovine serum for 60□ minutes (all Sigma-Aldrich), followed by overnight incubation with antibody and secondary antibody staining. Images were collected using ×40 (NA 1.25) oil immersion objectives on FV3000 confocal microscope (Olympus, Tokyo, Japan) with Olympus software and ImageJ software (NIH). Images were 3D-reconstructed using Volocity Image Analysis software (Improvision, Coventry, England). Primary antibody mouse anti-MUC2 (Santa Cruz Biotechnology, Dallas, TX; sc7314) at 1:100 dilutions was used.

#### ASMase activity

The enzymatic hydrolysis of SM to ceramide and phosphorylcholine by acid sphingomyelinase was measured with the Acid Sphingomyelinase Activity Assay Kit (Echelon Biosciences, K-3200) according to the manufacturer’s instructions. Caco2/BBe cells and colonoids grown in Transwell inserts were washed with cold PBS and then scraped in 1 mM PMSF on ice. Subsequently, cells were sonicated in an ice water bath sonicator for 10 min and then were disrupted by three rounds of freeze-thaw cycles with liquid nitrogen with vortexing in between each cycle. Extracts were clarified by centrifugation for 12 min at 16,000g at 4 C, and protein concentrations were determined by the BCA protein assay (Thermo Scientific). The enzymatic assay was carried out in 96-well plates. Each well contained 5 μg of protein in 20 μl of 1 mM PMSF. ASM standards were prepared in 1 mM PMSF as well. Substrate buffer (Echelon Biosciences, K-3203), 30 μl, was added to each well containing 20 μl of sample or standard. Then 50 μl/well of the diluted substrate (Echelon Biosciences, K-3202) in substrate buffer was added to each well. The plate was incubated for 3 h at 37 C with agitation. Stop buffer (Echelon Biosciences, K3204) was added to each well, and fluorescence was measured after 10 min of shaking at RT using FLUOstar Omega plate reader at 360 nm excitation and 460 nm emission. Data are presented as pmol substrate degraded/μg protein/min.

#### Immunoblotting

Cells were rinsed 3 times and harvested in phosphate-buffered saline by scraping. Cell pellets were collected by centrifugation, solubilized in lysis buffer (60 mmol/L HEPES, 150 mmol/L NaCl, 3 mmol/L KCl, 5 mmol/L EDTA trisodium, 3 mmol/L ethylene glycol-bis(β-aminoethyl ether)- *N,N,N’,N’* -tetraacetic acid, 1 mmol/L Na _3_ PO _4_, and 1% Triton X-100, pH 7.4) containing a protease inhibitor cocktail, and homogenized by sonication. Protein concentration was measured using the bicinchoninic acid method. Proteins were incubated with sodium dodecyl sulfate buffer (5 mmol/L Tris-HCl, 1% sodium dodecyl sulfate, 10% glycerol, 1% 2-mercaptoethanol, pH 6.8) at 37°C for 10 minutes, separated by sodium dodecyl sulfate–polyacrylamide gel electrophoresis on a 10% acrylamide gel, and transferred onto a nitrocellulose membrane. The blot was blocked with 5% nonfat milk, and probed with primary antibodies against DRA (mouse monoclonal, 1:500, sc-376187; Santa Cruz), CFTR antibody 217 (1:200 dilution; the University of North Carolina at Chapel Hill, Chapel Hill, NC, and Cystic Fibrosis Foundation Therapeutics, Inc.), glyceraldehyde-3-phosphate dehydrogenase (mouse monoclonal, 1:5000, G8795; Sigma-Aldrich), β-actin (mouse monoclonal, 1:5000, A2228; Sigma-Aldrich) overnight at 4°C, followed by secondary antibody against mouse IgG (1:10,000) for 1 hour at room temperature. Protein bands were visualized and quantitated using an Odyssey system and Image Studio software (LI-COR Biosciences, Lincoln, NE).

#### Surface Biotinylation

At 4°C, cells were incubated with 1.5 mg/mL N-hydroxysulfosuccinimide (NHS)-SS-biotin N-Hydroxysulfosuccinimide- and solubilized by lysis buffer. A small proportion (50μg) of the protein lysate was collected as the total lysate, while the rest was incubated with avidin-agarose beads overnight. The beads were centrifuged and washed with lysis buffer containing 0.1% Triton X-100. Biotinylated proteins were eluted from the beads and collected as the surface fraction. Immunoblotting was performed as described earlier and the percentage of surface expression of DRA was calculated as previously reported by loading equal volumes for each total and surface fraction^21^. The surface-to-total ratios were calculated separately for the DRA upper band (highly glycosylated, ~102 kilodaltons in size), the lower band (core glycosylated, ~85 kilodaltons in size), as well as for both bands together.

#### Measurement of Cl^-^/HCO _3_^-^ Exchange Activity

A detailed procedure for measuring Cl^-^ /HCO_3_^-^ exchange activity in Caco-2/BBe monolayers has been described previously^21^. In brief, the activity was fluorometrically measured by using the pHi-sensitive dye BCECF-AM in a customized chamber allowing simultaneous but separate apical and basolateral superfusion. The monolayers were rinsed and equilibrated in sodium solution (138 mM NaCl, 5 mM KCl, 2 mM CaCl2, 1 mM MgSO4, 1 mM NaH2PO4, 10 mM glucose, 20 mM HEPES, pH 7.4) for 60 minutes at 37 °C, then loaded with BCECF-AM (10 mM) in the same solution for another 30 min, and mounted in a fluorometer (Photon Technology International, Birmingham, NJ). The basolateral surface was superfused continuously with Cl-solution (110 mM NaCl, 5 mM KCl, 1 mM CaCl_2_, 1 mM MgSO_4_, 10 mM glucose, 25 mM NaHCO_3_, 1 mM amiloride, 5 mM HEPES, 95% O_2_/5% CO_2_), while the apical side was superfused with a shift between Cl-solution or Cl^-^ -free solution (110 mM Na-gluconate, 5 mM K-gluconate,5 mM Ca-gluconate, 1 mM Mg-gluconate, 10 mM glucose, 25 mM NaHCO_3_, 1 mM amiloride, 5 mM HEPES, 95% O_2_/5% CO_2_). The switch between Cl^-^ solution and Cl^-^ -free solution results in HCO_3_^-^ uptake across the apical membrane mediated by Cl^-^ /HCO_3_^-^exchangers (such as DRA). The subsequent change of intracellular pH was recorded and the rate of initial alkalization was calculated using Origin 8.0 (Origin-Lab, Northampton, MA) as an indicator of Cl^-^ /HCO_3_^-^ exchange activity. The specificity of the contribution of DRA to the rate of alkalinization was determined during standardization by determining the prevention of alkalinization by the apical presence of the DRA inhibitor Ao250 (10μM) (gift of A. Verkman).

#### Detergent-soluble (DS) and -insoluble (DI) Fractions of Total Membranes

Separation of the total membrane preparation into detergent-soluble and detergent-insoluble fractions was as we reported previously^7, 47^. Post-confluent Caco-2/BBe cells in submerged culture or after ALI modifications were homogenized by passing them through a 1-ml syringe/26-gauge needle in TNE buffer A containing 25 mM Tris, pH 7.4, 150 mM NaCl, 50 mM NaF, 5 mM EDTA, 1 mM Na_3_VO_4_, and protease inhibitors. Nuclei and debris were removed by centrifugation at 3000 × *g* for 15 min at 4 °C. The total membranes were pelleted by ultracentrifugation at 100,000 × *g* for 30 min at 4 °C. Total membranes were then solubilized with cold buffer A supplemented with 0.5% Triton X-100 and then incubated at 4 °C for 30 min on a rotary shaker and subjected to ultracentrifugation at 100,000 × *g* for 30 min at 4 °C. The supernatant is referred to as the DS fraction. The pellet was resuspended in 1× SDS-PAGE loading buffer (equal volume to the DS fraction) as the DI fraction.

#### Reverse transcription and real-time PCR

A reported procedure to measure the transcriptional expression of target genes using primer pairs was followed^25, 44^. In brief, the PureLink RNA Mini Kit (Life Technologies) was used to extract total RNA from Caco-2/BBe cells or colonoid monolayers. Then the complementary DNA was synthesized from the extracted RNA using SuperScript VILO Master Mix (Life Technologies). Quantitative real-time PCR (qRT-PCR) was performed on a QuantStudio 12K Flex real-time PCR system (Applied Biosystems, Foster City, CA) by using Power SYBR Green Master Mix (Life Technologies). Samples were run in triplicate, and the relative mRNA expression level of targeted genes was calculated from the 2-ΔΔCT method normalized with the vehicle control and housekeeping β-actin RNA.

#### Measurement of active anion secretion (short-circuit current, Isc)

The short-circuit current as an indicator of active electrogenic anion secretion in Caco-2/BBe cells was determined by the Ussing chamber/Voltage Clamp Technique^21^. In brief, Caco-2/BBe monolayers were mounted in Ussing chambers and incubated in Krebs-Ringer bicarbonate (KBR) buffer (115 mM NaCl, 25 mM NaHCO_3_, 0.4 mM KH_2_PO_4_, 2.4 mM K_2_HPO_4_, 1.2 mM CaCl_2_, 1.2 mM MgCl_2_, pH 7.4) continuously gassed with 95% O_2_/5% CO_2_ at 37°C and connected to a voltage-current clamp apparatus (Physiological Instruments) via Ag/AgCl electrodes and 3 M KCl agar bridges. 10 mM glucose was supplemented as an energy substrate on the basolateral side, while 10 mM mannitol was added to the apical chamber to maintain the osmotic balance. Current clamping was employed and short-circuit current was recorded every 1 or 5 seconds by the Acquire & Analyze software 2.2.2 (Physiological Instruments, San Diego, CA, USA). 10 μM forskolin was added to the apical and basal chamber to initiate cAMP-stimulated anion secretion.

#### Statistics

GraphPad Prism (version 6.01, GraphPad Software, San Diego, CA) was used to perform the statistical analysis. Data are mean ± s.e.m. of at least three independent experiments, with an error bar equaling one s.e.m. A two-tail Student’s t-test was used for statistical comparison between two groups, while a one-way analysis of variance (ANOVA) followed by post hoc Turkey was adopted if more than two different groups were compared. P < 0.05 was considered as the threshold of statistical significance.

All authors had access to the study data and reviewed and approved the final manuscript.

## Abbreviations

ASMase: Acid Sphingomyelinases
SLC26A3: solute carrier family 26 member 3

## Acknowledgments

We thank the Hopkins Conte Basic and Translational Digestive Diseases Research Core Center Integrated Physiology and Translational Research Enhancement Cores for their contributions to obtaining and maintaining colonoids.

## Patient consent

Obtained.

## Ethics approval

Johns Hopkins Medicine IRB committee.

