## Supplemental figure for "Air-Liquid-Interface Reorganizes Membrane Lipid and Enhance Recruitment of Slc26a3 to Lipid-Rich Domains in Human Colon"

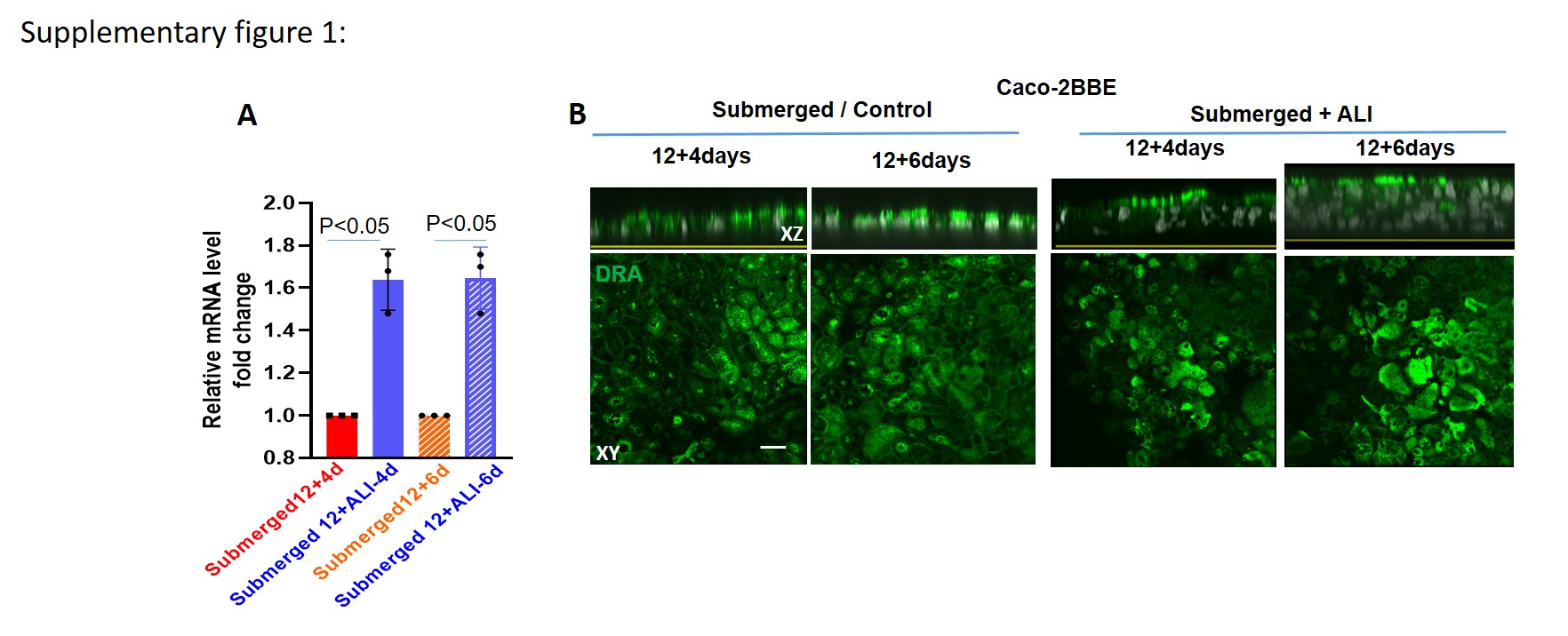

**Supplementary Figure 1: Confluent Caco-2/BBe monolayer changes to multilayers when exposed to ALI culture for more than 2 days.** **(A)** DRA mRNA expression changed between the submerged culture and after ALI exposure for the indicated number of days. Relative mRNA expression of DRA by quantitative PCR. Messenger RNA levels are normalized to 18S ribosomal RNA expression. The result is normalized to the control set as 1 and expressed as fold change. Results are Means ± SEM of 3-5 independent experiments. P values represent unpaired Student’s t-test. **(B)** Confocal fluorescence microscopy of DRA (green) expression and nucleus stained with DAPI (blue) under different conditions; top: XZ viewing direction; below: single plane of Z-stack section at the level of the apical membrane is shown. Scale bars10 μm.
